# Discovery of potent inhibitors of PL^pro^CoV2 by screening a library of selenium-containing compounds

**DOI:** 10.1101/2020.05.20.107052

**Authors:** Ewelina Węglarz-Tomczak, Jakub M. Tomczak, Mirosław Giurg, Małgorzata Burda-Grabowska, Stanley Brul

## Abstract

A collection of twelve organoselenium compounds, structural analogues of antioxidant drug ebselen were screened for inhibition of the papain-like protease (PL^pro^) from the acute respiratory syndrome coronavirus 2 (SARS-CoV-2, CoV2). This cysteine protease, being responsible for the hydrolysis of peptide bonds between specific amino acids, plays a critical role in CoV2 replication and in assembly of new viral particles within human cells. The activity of the PL^pro^ CoV2 is essential for the progression of coronavirus disease 2019 (COVID-19) and it constitutes a key target for the development of anti-COVID-19 drugs. Here, we identified four strong inhibitors that bind favorably to the PL^pro^ CoV2 with the *IC*_50_ in the nanomolar range.

## Introduction

Severe acute respiratory syndrome coronavirus 2 (SARS-CoV-2) is the cause of the coronavirus disease 2019 (COVID-19) in the coronavirus pandemic that on May 19, 2020 affected around 4.5 million people in over 188 countries [1]. The scale of the pandemic, easiness of its spreading [2] and variety of potential complications [3], which yet remain not fully understood, position COVID-19 among serious diseases faced by humankind so far.

The extensive research on developing new antiviral drugs for COVID-19 led to identifying two potential targets being cysteine proteases that play a vital role in viral replication. Main protease (M^pro^, also known as chymotrypsin-like protease 3CLpro) is responsible for polypeptide processing during virus replication [4,5]. Papain-like protease (PL^pro^) additionally to its role in virus replication assembles new viral particles within human cells [6,7]. Harcourt et al. [6] showed that PL^pro^ is a key enzyme in the pathogenesis of SARS-CoV-1 [8], the causative agent of the fatal global outbreak of respiratory disease in humans during 2002–2003 [9]. In the study presented by Shin et al. [10], PL^pro^ from novel SARS-CoV-2 is claimed to be the essential viral enzyme that weakens the antiviral immune response and helps to take advantage of the host’s immune system for its own benefit. Thus, current understanding of mechanisms of SARS-CoV-2 indicates that blocking PL^pro^ seems to be crucial for further stopping virus spread.

The identification of SARS-CoV-2 PL^pro^ as an essential viral enzyme [10] offers a great possibility for drug discovery. In recent studies, peptide analogues [11] and ebselen [12] have been identified as highly active inhibitors for PL^pro^. Ebselen is a seleno-organic drug with well-known anti-inflammatory, anti-atherosclerotic, cytoprotective properties and low toxicity in humans [13]. It has been proven to be an effective therapeutic agent in multiple diseases like cancer and hepatitis C virus [14-18].

In this paper, we present inhibitory activity of twelve ebselen derivatives obtained by substitution/functionalization of the phenyl ring towards SARS-CoV-1 PL^pro^ (PL^pro^SARS) and SARS-CoV-2 PL^pro^ (PL^pro^CoV2). Previously, these derivatives were proven to be highly effective towards human methionine aminopeptidase 2 [19] and antiviral and antimicrobial agents [20]. We show that some of them indeed possess higher activity than ebselen, that has been recently reported as PL^pro^CoV2 inhibitor [12], and, thus, could be considered as novel potential drugs against COVID-19.

## Results

Twelve ebselen derivatives/analogues compounds, seven benzisoselenazol-3(2*H*)-ones (**1a**–**g**) and four 2,2′-dicarbamoyldiaryl diselenides (**2a**-**2f**) were employed for inhibitory studies toward PL^pro^ from SARS and CoV2. The first group included 2-phenylbenzisoselenazol-3(2*H*)-ones with the phenyl ring monosubstituted with functional group, such as Me (**1c**), OH (**1d**), OMe(**1e**) and analogues of ebselen based on the benzisoselenazol-3(2*H*)-one core modified in position 2 (the nitrogen atom) by adding -CH_2_-(**1f**) and -CH_2_CH_2_-(**1g**) (compound **1f** is additional monosubstituted by t-Bu). The second group constitutes acyclic diselenide form of ebselen or its derivatives (**2d** and **2e**) containing two atoms of selenium per molecule.

Organoselenium compounds were found to be irresistible inhibitors toward both enzymes. Similar to recently published ebselen [12] they show irreversibility of the mode of action. All of phenylbenzisoselenazol-3(2*H*)-ones inactivated completely PL^pro^Cov2 in concentration equal 20μM (see Table 1). In the case of PL^pro^SARS the range of inhibition was from 50% to 100% (see Table 1).

**Table 1.**
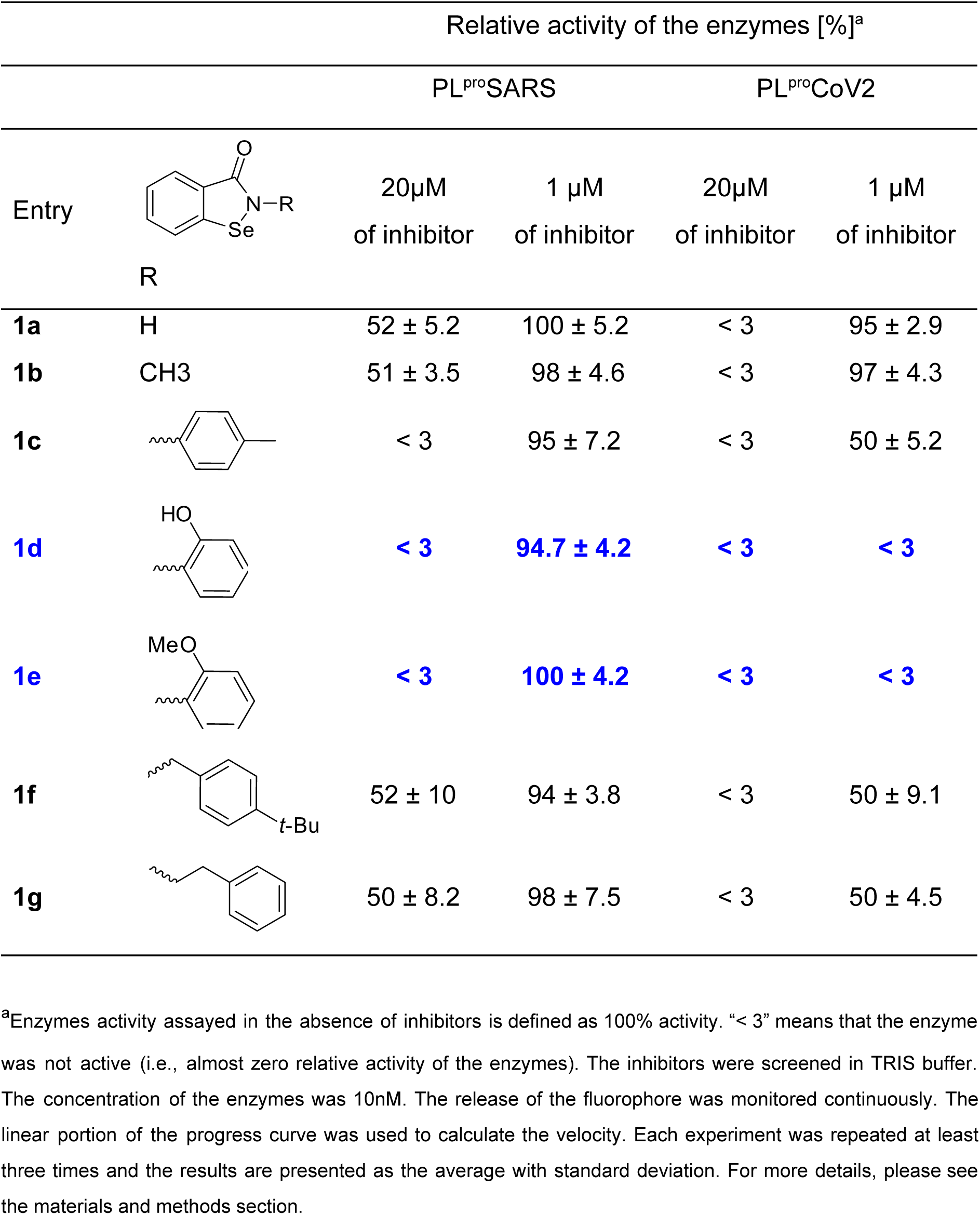
Inhibitory activity of ebselen derivatives obtained by substitution/functionalization of the phenyl ring towards PL^pro^SARS and PL^pro^CoV2. The most significant inhibition is highlighted in blue.

| Relative activity of the enzymes [%]<sup>a</sup> |  |  |  |  |  |
| --- | --- | --- | --- | --- | --- |
|  |  | PL<sup>pro</sup>SARS |  | PL<sup>pro</sup>CoV2 |  |
| Entry | R | 20μM<br>of inhibitor | 1 μM<br>of inhibitor | 20μM<br>of inhibitor | 1 μM<br>of inhibitor |
| **1a** | H | 52 ± 5.2 | 100 ± 5.2 | < 3 | 95 ± 2.9 |
| **1b** | CH<sub>3</sub> | 51 ± 3.5 | 98 ± 4.6 | < 3 | 97 ± 4.3 |
| **1c** |  | < 3 | 95 ± 7.2 | < 3 | 50 ± 5.2 |
| **1d** |  | < 3 | 94.7 ± 4.2 | < 3 | < 3 |
| **1e** |  | < 3 | 100 ± 4.2 | < 3 | < 3 |
| **1f** |  | 52 ± 10 | 94 ± 3.8 | < 3 | 50 ± 9.1 |
| **1g** |  | 50 ± 8.2 | 98 ± 7.5 | < 3 | 50 ± 4.5 |

<sup>a</sup>Enzymes activity assayed in the absence of inhibitors is defined as 100% activity. “< 3” means that the enzyme was not active (i.e., almost zero relative activity of the enzymes). The inhibitors were screened in TRIS buffer. The concentration of the enzymes was 10nM. The release of the fluorophore was monitored continuously. The linear portion of the progress curve was used to calculate the velocity. Each experiment was repeated at least three times and the results are presented as the average with standard deviation. For more details, please see the materials and methods section.

The experiment with 1μM of the inhibitor led to identify highly active ligands. Two derivatives of ebselen with substitution by hydroxy (**1c**) or methoxy group (**1d**) inhibited CoV2 in 100% in this condition. Affinity of diselenide orthologs (**2c** and **2d**) were overall in the same range (Table 1 and 2). Whereas, replacing phenyl in position 2 with less hydrophobic substituents, hydrogen (**1a**) or methyl (**1b**), was not beneficial, similar to substitution by methyl in the *para* position (**1b**). We observed a similar relationship for diselenide (Table 1). All compounds were less active toward PL^pro^SARS. Only compounds that are substituted derivatives of ebselen and their diselenide orthologs showed significant inhibition in concentration 20 μM.

**Table 2.**
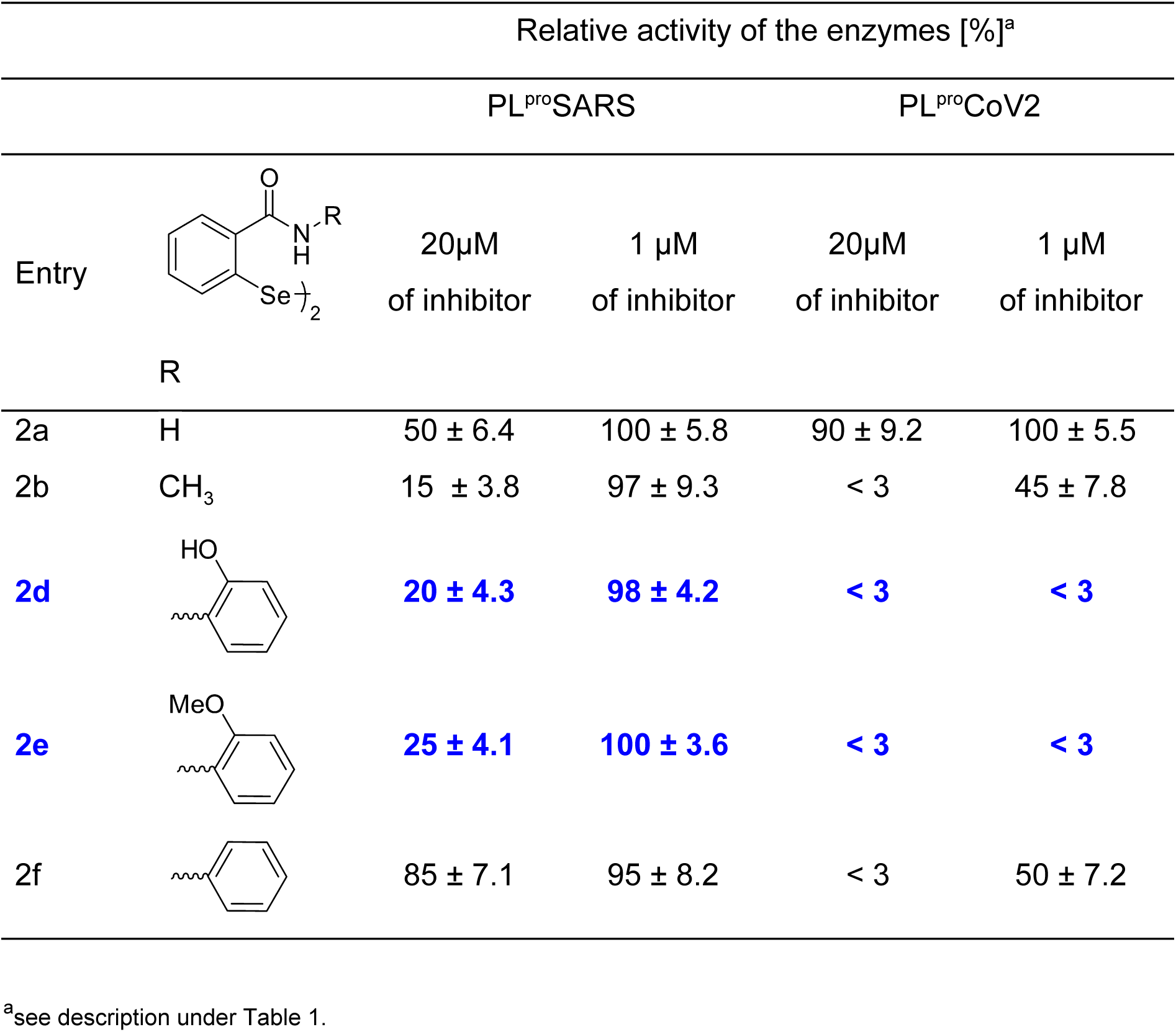
Inhibitory activity for diselenides, the acyclic dimeric forms of ebselen and analogues, toward PL^pro^SARS and PL^pro^CoV2.

| Relative activity of the enzymes [%]<sup>a</sup> |  |  |  |  |  |
| --- | --- | --- | --- | --- | --- |
|  |  | PL<sup>pro</sup>SARS |  | PL<sup>pro</sup>CoV2 |  |
| Entry | R | 20μM<br>of inhibitor | 1 μM<br>of inhibitor | 20μM<br>of inhibitor | 1 μM<br>of inhibitor |
| 2a | H | 50 ± 6.4 | 100 ± 5.8 | 90 ± 9.2 | 100 ± 5.5 |
| 2b | CH<sub>3</sub> | 15 ± 3.8 | 97 ± 9.3 | < 3 | 45 ± 7.8 |
| **2d** |  | **20 ± 4.3** | **98 ± 4.2** | **< 3** | **< 3** |
| **2e** |  | **25 ± 4.1** | **100 ± 3.6** | **< 3** | **< 3** |
| 2f |  | 85 ± 7.1 | 95 ± 8.2 | < 3 | 50 ± 7.2 |

<sup>a</sup>see description under Table 1.

The inhibitory potency was further investigated for the most significant ligands with PL^pro^CoV2 and we found *IC*_50_ value in nanomolar range for **1c, 1d, 2c** and **2d** (Table 3). All four compounds appeared to be very effective inhibitors of PL^pro^ from CoV2, with the *IC*_50_ constants in the nanomolar range, e.g. 236 nM for compound **1e** with methyl substitution in the *para* position.

**Table 3.**
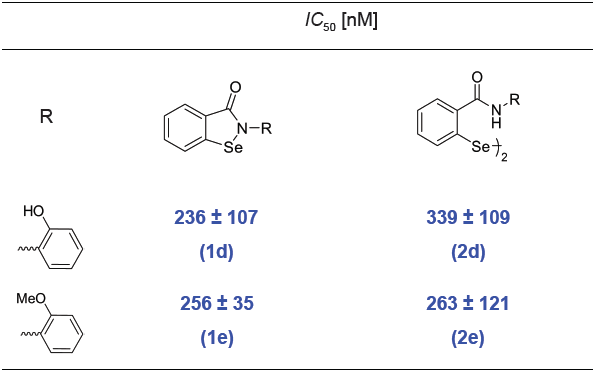
Inhibitory activity for compounds **1d, 1e** (ebselen derivatives obtained by substitution of the phenyl ring) and **2e** and **2f** (diselenides, the acyclic forms of **1d** and **1e**).

| $IC_{50}$ [nM] | | |
| --- | --- | --- |
| R |  |  |
|  | **236 ± 107**<br>**(1d)** | **339 ± 109**<br>**(2d)** |
|  | **256 ± 35**<br>**(1e)** | **263 ± 121**<br>**(2e)** |

The most significant results obtained in this study were further illustrated with molecular modeling (Figure 1 and 2). The modeled interactions do not show significant changes in the overall binding mode architecture compared with the ebselen-PL^pro^CoV2 [12]. Similar to ebselen complexes with enzyme, hydroxyl derivative occupies the same intersection between catalytic Cys111–His272–Asp286 triad and Trp106 and it is wrapped by other Tyr268, Ala289 and Leu298 making with them face-to-edge stacking and π-alkyl interactions, respectively (Figure 1). Additionally, *Se*-phenyl and an indole from His272 forms π–π stacking interactions. In the case of compound **1d** possessing hydroxyl group, negatively charged oxygen atoms coordinate the carboxyl group from Asp286 (Figure 1). Whereas the methoxy group in compound **1e** forms π-alkyl interactions with aromatic rings from His272 and Tyr268 (Figure 2).

**Figure 1.**
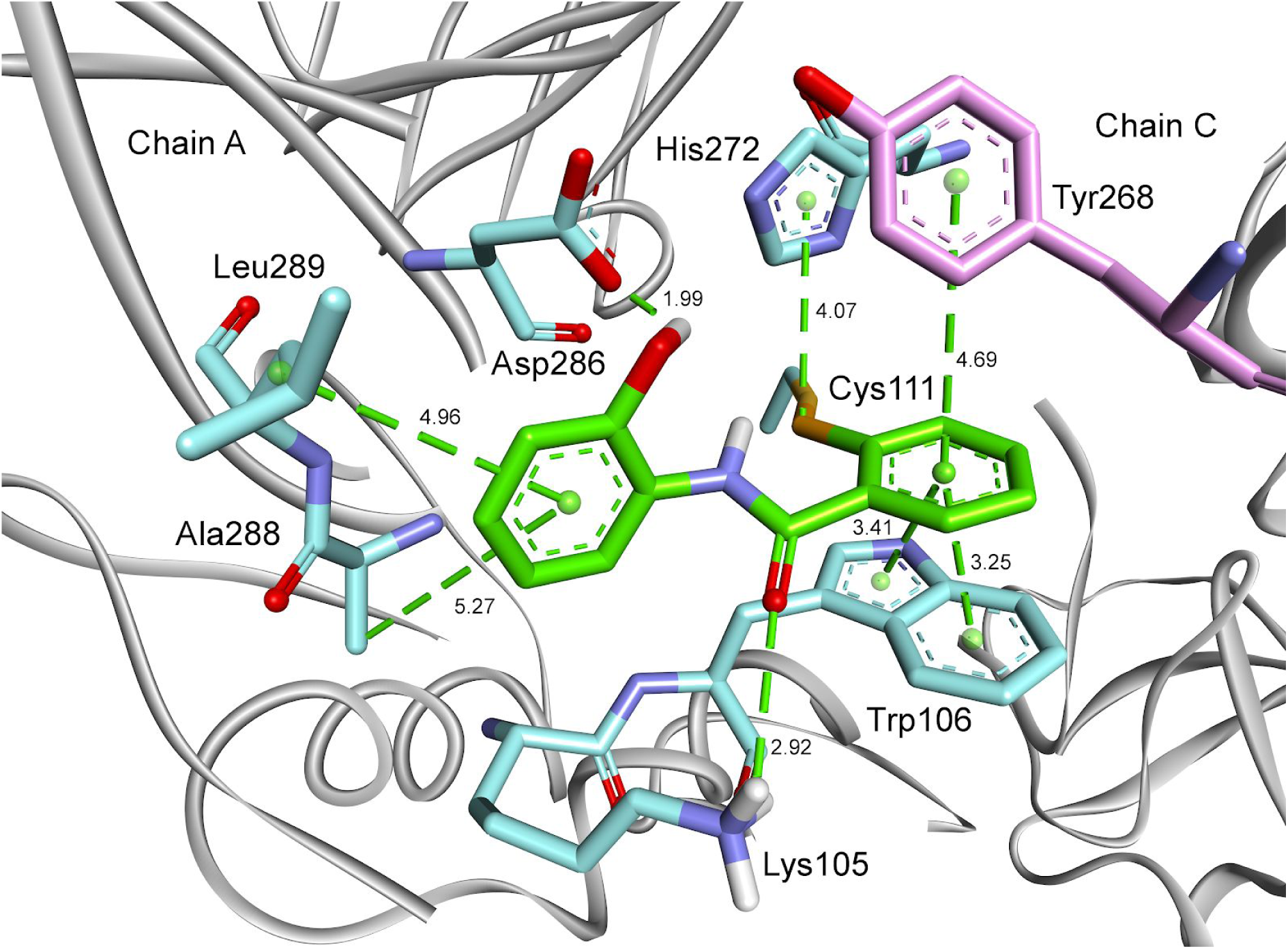
A model of the complex of compound **1d** with SARS-CoV-2 PL^pro^ (PDB: 6W9C [22]).

**Figure 2.**
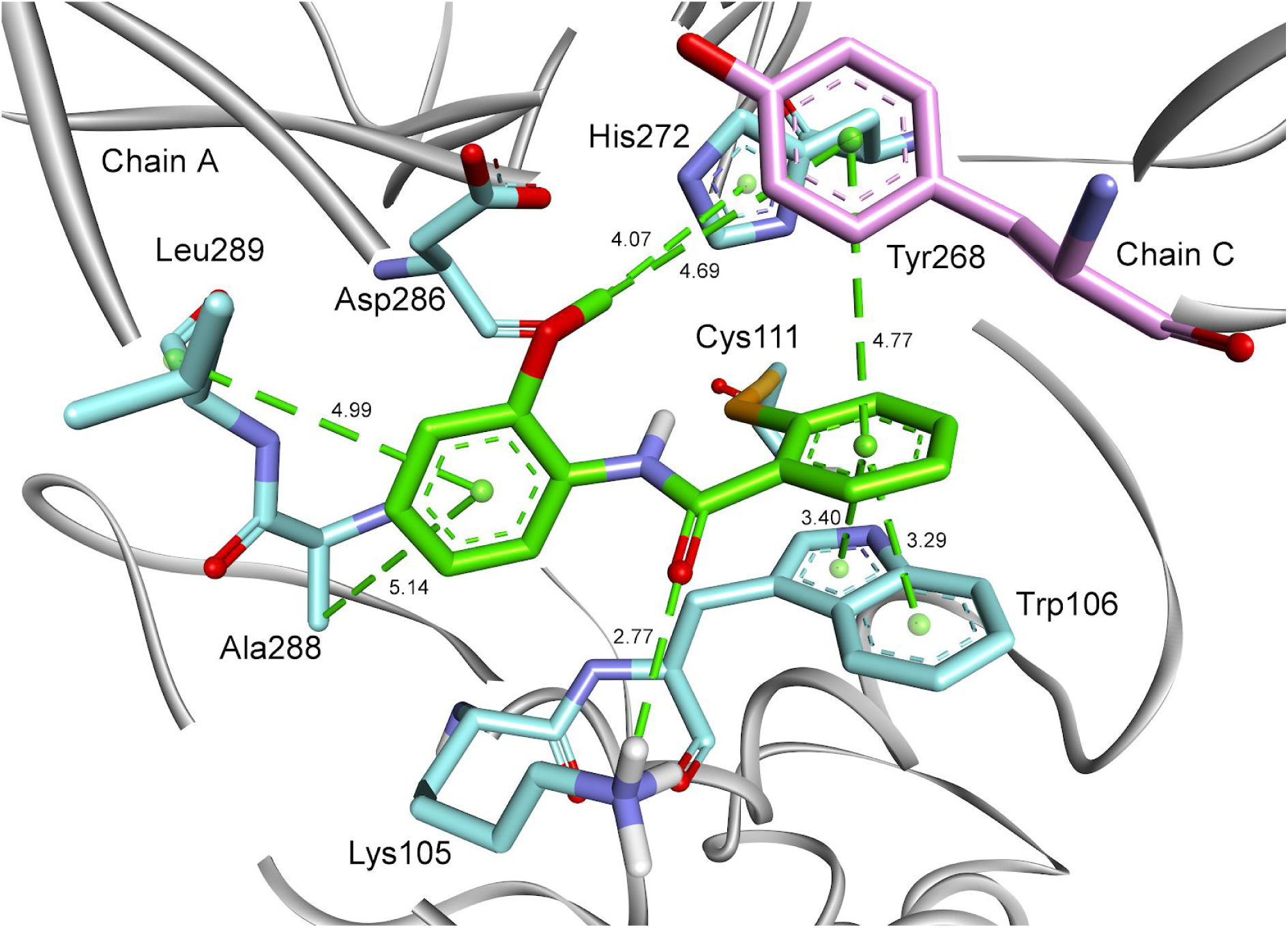
A model of the complex of compound **1e** with SARS-CoV-2 PL^pro^ (PDB: 6W9C [22]).

## Discussion

Inhibition of papain like protease from CoV2 has been recently identified as a potential approach to therapy of COVID-19 [10]. In this work, we used the ebselen derivatives/analogues library and performed a comprehensive inhibition study of PL^pro^CoV2. All of the tested compounds proved to be covalent inhibitors of the enzyme. Interestingly, all derivatives except **1a** blocked completely PL^pro^CoV2 at a high concentration of the inhibitor (20μM), but only some of them (**1c, 1d, 1e** and **2d, 2e**) inhibited PL^pro^SARS while rest of them failed. This outcome is even more apparent for the concentration of 20μM of an inhibitor. In the case of PL^pro^SARS, none of the presented ebselen derivatives was able to block the enzyme. However, **1d, 1e, 2d** and **2e** inhibited PL^pro^CoV2 at this concentration. The investigation of *IC*_50_ revealed that **1d, 1e, 2d** and **2e** obtained approximately 250 nM that is one magnitude better than in the case of ebselen (around 2 μM as reported in [12]).

In conclusion, we identified very effective inhibitors of PL^pro^CoV2, with the *IC*_50_ constants in the nanomolar range. Our findings provide evidence that ebselen derivatives with an additional hydroxy or methoxy group serve as highly potential prospective drugs against COVID-19.

## Material and methods

### 1. General

SARS-CoV-1 PLpro, SARS-CoV-2 PLpro and Ubiquitin-AMC were purchased as 32, 11 and 3 μM solutions, respectively, from R&D Systems.

All compounds were obtained and fully characterized in previous studies [19-21]. Their purity and homogeneity were confirmed by HRMS and _77_Se NMR.

### 2. Enzymes assay

The enzymes were dissolved in a 50 mM Tris-HCl buffer containing DTT (2 mM), NaCl (100mM) and 0.1 mg/mL albumin, at pH 7.5, and preincubated 30 min. Spectrofluorimetric measurements were performed in a 96-well plate format working at two wavelengths: excitation at 355 nm and emission at 460 nm. The release of the fluorophore was monitored continuously at the enzyme concentration of 10 nM. The linear portion of the progress curve was used to calculate velocity of hydrolysis.

### 3. Inhibition assay

The inhibitor was screened against recombinant PL^pro^SARS and PL^pro^CoV2 at 37°C in the assay buffer as described above. For steady state measurement the enzymes were incubated for 60min at 37°C with an inhibitor before adding the substrate to the wells. Eight different inhibitor concentrations were used. Value of the concentration of the inhibitor that achieved 50% inhibition (*IC*_50_) was taken from the dependence of the hydrolysis velocity on the logarithm of the inhibitor concentration [I].

### 4. Molecular Modeling

Molecular modeling studies were performed using the Discovery Studio 2020 (Dassault Systemes BIOVIA Corp). The crystal structure of the SARS-CoV-2 (PDB ID 6W9C [22]) with protons added (assuming the protonation state of pH 7.5) was used as the starting point for calculations of the enzyme complexed with ebselen. The partial charges of all atoms were computed using the Momany-Rone algorithm. Minimization was performed using the Smart Minimizer algorithm and the CHARMm force field up to an energy change of 0.0 or RMS gradient of 0.01. Generalized Born model was applied. The nonbond radius was set to 14 Å.

## Acknowledgments

We gratefully acknowledge the Dassault Systemes for the free license for BIOVIA: Discovery Studio package given for our research.

EWT is co-financed by a grant Mobilność Plus V from the Polish Ministry of Science and Higher Education (Grant No. 1639/MOB/V/2017/0).

## Author contributions

EWT conceived the project. EWT designed the research and experiments with contributions from JT and SB. Experimental work was done by EWT. Molecular docking was done by JT and EWT. MG and MB contributed to synthesis. EWT, JT and SB drafted and revised the manuscript.

## Competing interests

The authors declare no competing interests.

## Materials and correspondence

Correspondence and requests for materials should be addressed to EWT.

## Notes

### Competing Interest Statement

The authors have declared no competing interest.

### Summary of Updates

A typo in title fixed. A list of authors updated.

## Literature

1. WHO Novel Coronavirus (COVID-19) pandemic https://www.who.int/emergencies/diseases/novel-coronavirus-2019

2. Sanche S. et al. High Contagiousness and Rapid Spread of Severe Acute Respiratory Syndrome Coronavirus 2. Emerg. Infect. Dis. 26. (2020). doi: 10.3201/eid2607.200282

3. “Interim Clinical Guidance for Management of Patients with Confirmed Coronavirus Disease (COVID-19)”. U.S. Centers for Disease Control and Prevention (CDC). 4 April 2020. Retrieved 11 April 2020.

4. Rismanbaf A. Potential Treatments for COVID-19; a Narrative Literature Review. Arch Acad Emerg Med. 8, e29 (2020).

5. Wu C. et al. Analysis of therapeutic targets for SARS-CoV-2 and discovery of potential drugs by computational methods. Acta Pharm Sin B. in Press. (2020). doi: 10.1016/j.apsb.2020.02.008

6. Harcourt B. H. et al. Identification of severe acute respiratory syndrome coronavirus replicase products and characterization of papain-like protease activity. J. Virol. 78, 13600–13612 (2004).

7. Báez-Santos M., St. John S. E. & Mesecar A. D. The SARS-coronavirus papain-like protease: Structure, function and inhibition by designed antiviral compounds. Antiviral Res. 115, 21–38 (2015).

8. Thiel, V., ed. Coronaviruses: Molecular and Cellular Biology (1st ed.). Caister Academic Press (2007).

9. Summary Table of SARS Cases by Country, 1 November 2002–7 August 2003. WHO: Geneva (August 15, 2003). http://www.who.int/csr/sars/country/2003_08_15/en/

10. Shin D. et al. Inhibition of papain-like protease PLpro blocks SARS-CoV-2 spread and promotes anti-viral immunity. Nature (under review) (2020). doi: 10.21203/rs.3.rs-27134/v1

11. Rut W. et al. Activity profiling and structures of inhibitor-bound SARS-CoV-2-PLpro protease provides a framework for anti-COVID-19 drug design. bioRxiv preprint (2020). doi: 10.1101/2020.04.29.068890

12. Węglarz-Tomczak E. et al., Ebselen as a highly active inhibitor of PLproCoV2, bioRxiv 2020.05.17.100768 (2020). doi: 10.1101/2020.05.17.100768

13. Schewe T. Molecular actions of ebselen—an antiinflammatory antioxidant. Gen. Pharmacol.-Vasc. S. 26, p. 1153–1169 (1995)

14. Chantadul V. et al. Ebselen as template for stabilization of A4V mutant dimer for motor neuron disease therapy. Commun Biol. 3, 1–10 (2020). doi: 10.1038/s42003-020-0826-3.

15. Hanavan P.D. et al. Ebselen inhibits QSOX1 enzymatic activity and suppresses invasion of pancreatic and renal cancer cell lines. Oncotarget 6 18418–18428 (2015).

16. Yan J. et al. Design, synthesis, and biological evaluation of benzoselenazole-stilbene hybrids as multi-target-directed anti-cancer agents. Eur. J. Med. Chem. 95. 220 (2015).

17. Mukherjee S. et al. Ebselen inhibits hepatitis C virus NS3 helicase binding to nucleic acid and prevents viral replication. ACS Chem. Biol., 9, 2393 (2014).

18. Garland, M. et al.. The Clinical Drug Ebselen Attenuates Inflammation and Promotes Microbiome Recovery in Mice after Antibiotic Treatment for CDI. Cell Rep. Med. 1, p.100005 (2020).

19. Węglarz-Tomczak, E., Burda-Grabowska, M., Giurg, M., & Mucha, A. Identification of methionine aminopeptidase 2 as a molecular target of the organoselenium drug ebselen and its derivatives/analogues: Synthesis, inhibitory activity and molecular modeling study. Bioorg. Med. Chem. Lett. 26, 5254–5259 (2016).

20. Pietka-Ottlik, M., et al. Synthesis of new alkylated and methoxylated analogues of ebselen with antiviral and antimicrobial properties. Arkivoc, 546–556 (2017).

21. Giurg M. et al. Reaction of bis[(2-chlorocarbonyl)phenyl] Diselenide with Phenols, Aminophenols, and Other Amines towards Diphenyl Diselenides with Antimicrobial and Antiviral Properties, Molecules 22, 974 (2017).

22. Osipiuk, J. et al. The crystal structure of papain-like protease of SARS CoV-2. doi: 10.2210/pdb6W9C/pdb

